# *In vitro* validation of neoantigen prediction algorithm for developing personalized cancer vaccine therapy

**DOI:** 10.1101/2021.12.12.472304

**Authors:** Yun-jeong Choe, Eunyoung Kim, Jooyeon Oh, Miran Jang, Weixan Fu, Hanna Lee, Minho Chung, Kyung-Ho Pyo, Chung-Bong Synn, Sora Kim, Yohan Yang, Ahyeon Kim, Byung Chul Cho, Han Sang Kim, Sangwoo Kim, Beatriz Carreno, Jee Ye Kim, Soonmyung Paik

**Author notes:** Co-first authors (Yun-jeong Choe, Eunyoung Kim).

## Abstract

**Background:** The development of personalized neoantigen-based therapeutic cancer vaccines relies on computational algorithm-based pipelines. One of the critical issues in the pipeline is obtaining higher positive predictive value (PPV) performance, i.e., how many are immunogenic when selecting the top 5 to 20 candidate neoepitopes for the vaccination. We attempted to test the PPV of a neoepitope prediction algorithm Neopepsee.

**Methods:** Six breast cancer patients and patient-derived xenografts from three lung cancer patients and their paired peripheral blood samples were subjected to whole-exome and RNA sequencing. Neoantigen was predicted using two different algorithms (Neopepsee and pVACseq). Response of induced memory T cells to neopeptide candidates was evaluated by IFN-γ Enzyme-linked immune absorbent spot (ELISpot) assays of peripheral blood mononuclear cell (PBMC) from three HLA-matched donors. Positive ELISpot response to a candidate peptide in at least 2 of 3 donor PBMC was regarded as an immunogenic response.

**Results:** Neopepsee predicted 159 HLA-A matched neoepitope candidates out of 898 somatic mutations in nine patients (six breast and three lung cancer patients), whereas pVACseq predicted 84 HLA-A matched candidates. A total of 26 neopeptide candidates overlapped between the two predicted candidate pools. Among the candidates, 28 (20%, 28/ 137) and 15 (20%, 15/ 75) were positive by ELISpot assay, respectively. Among 26 overlapped candidates, 20 could be tested, and 7 of them (35%) were validated by ELISpot. Neopepsee identified at least one neoepitope in 7 of 9 patients (range 0-6), compared to 6 by pVACseq (range 0-5).

**Conclusion:** As suggested by Tumor Neoantigen Selection Alliance (TESLA), our results demonstrate low PPV of individual prediction models as well as the complementary nature of the Neopepsee and pVACseq and may help design neoepitope targeted cancer vaccines. Our data contribute a significant addition to the database of tested neoepitope candidates that can be utilized to further train and improve the prediction algorithms.

## INTRODUCTION

Cancer results from the accumulation of somatic mutations.^1^ The Cancer Genome Atlas (TCGA) result demonstrated that most solid tumor cells have an average of 50 non-synonymous mutations.^2^ Minor fractions of peptides encoded by the somatic mutations in cancer cells are presented by major histocompatibility complex (MHC) class I proteins and induce an anti-tumor immune response by T lymphocytes.^3^Such mutant peptides are called “neoepitopes” or “neoantigens”.

Neoepitopes are thought to be the best targets for therapeutic cancer vaccines since, unlike non-mutated tumor-associated antigens, they are expected to induce robust T-cell immune response due to lack of central tolerance and are not likely to induce autoimmunity to normal cells.^4–6^ Adoptive transfer of *ex vivo* expanded tumor-infiltrating lymphocytes enriched for neoepitope specific T-cells resulted in remarkable responses in some patients, especially when administered together with immune checkpoint inhibitors, demonstrating that mutated neoepitopes are valid therapeutic targets.^7^

Early results from a dendritic cell, mRNA, or peptide vaccination against neoepitopes demonstrated induction of anti-tumor immunity and evidence of epitope spreading as well as a suggestion of clinical efficacy when used in combination with immune checkpoint inhibitors.^8–11^ Near absence of dose limiting-toxicities due to off-target immune response against normal tissue in both therapeutic vaccine and adoptive cell transfer trials is reassuring.^8–11^ With the advancement of timely identification of neopeptide candidates, many clinical trials are now ongoing with a variety of vaccine platforms targeting neoepitopes.^4,5,12^

Due to the time constraint in caring for patients diagnosed with advanced cancer, it is difficult to experimentally validate each candidate neoepitopes *in vitro* before administering the vaccine to patients. Therefore, most neoepitope vaccine trials rely on computational prediction algorithms to select candidate neoepitopes in personalized vaccines. In reported clinical trials, the immune response was observed for less than 50% of the vaccinated peptides or mRNA encoding the peptides.^8–11^ On the other hand, a study suggests the existence of the “Inhibigens”, neoepitopes that induce immune suppression rather than stimulation.^13^ In preclinical models, adding an inhibigen could prevent induction of therapeutic immune response by immune-stimulating neoepitopes.^13^ Therefore, the accuracy of the computational prediction algorithm is important when designing a vaccine with predicted neoepitope candidates without in vitro validation of their T-cell immunogenicity. Of note, the positive predictive value (PPV) is one of the most critical measurements in designing a vaccine, i.e., how many are true positives when selecting the top 5 to 20 candidate neoepitopes for the vaccination. A recent report from the Tumor Neoantigen Selection Alliance (TESLA) showed that PPV is only 7% on average across 25 participating institutions.^14^ PPV was improved up to 50% when an ensemble of different prediction algorithms was combined.

Previously, we reported a neoepitope prediction model, “Neopepsee”, which was trained with multiple features that may influence the immunogenicity of mutated peptides.^15^Although Neopepsee showed good performance compared to previously reported algorithms in a validation set *in silico*, experimental validation was lacking. We have conducted a prospective experimental validation of Neopepsee in 9 patients with breast and lung cancer using ELIspot assay with HLA matched donor peripheral blood mononuclear cells (PBMC).^16,17^ As a comparator, we used pVacSeq, a widely used prediction algorithm.^18^ As a corollary to the study, we examined whether whole-exome and RNAseq-based HLA typing are reliable by comparing the results with two-digit clinical HLA genotyping and dedicated NGS-based HLA typing kit.

## METHODS

### Subjects and ethics requirements

Fresh tumor and peripheral blood samples were obtained from six breast cancer patients undergoing surgery at Yonsei Cancer Center with written informed consent and institutional review board approval (IRB approval number:4-2017-0715). All patients except one had not received any systemic treatment or radiotherapy in the biopsied area before surgery.

Anonymized patient-derived xenografts (F4 generation) with paired peripheral blood samples were available for three lung cancer patients with informed consent (IRB approval number 4-2016-0788). Leukoreduction system (LRS) chamber blood cells from 50 healthy donors were obtained from the central blood bank of Korea (IRB approval number: 4-2018-0803).

### HLA genotyping

We used Sanger sequencing-based clinical HLA genotyping results with two-digit resolution (BIOWITHUS Inc. Seoul, Korea). In addition, we applied Optitype ^19^ and HLAminer ^20^and used the Omixon Holotype HLA™ kit (Omixon Biocomputing Ltd, Budapest, Hungary) to evaluate NGS-based HLA typing methods.

### Neoepitope candidate selection

Paired whole-exome sequencing (WES) and RNA sequencing were performed at Yonsei Genome Center. WES library was generated using SureSelect Exome V7 (Agilent Technologies, CA, USA) for breast cancer samples or V5 for lung cancer samples, and sequencing data was produced using Novaseq6000 (Illumina, CA, USA). Multiplexing for sequencing was designed to acquire more than 200X depth for tumor samples and 100X depth for matched normal samples, respectively. Total RNA sequencing library was generated using TruSeq stranded total RNA library prep kit (Illumina), and sequencing data was produced using Novaseq 6000 (except for sample Neo 1, which was sequenced using Nextseq550) to acquire more than 5 G data for each sample.

Tumor-normal paired WES reads were aligned to the human reference genome with BWA-MEM.^21^ To reduce mouse contamination from patient-derive xenografts (PDX) sample sequencing data, we made a concatenated reference of mouse and human as recommended and performed an alignment with concatRef on PDX of 3 lung cancer patients.^22^ We used Picard (version 2.19) (https://broadinstitute.github.io/picard/) to sort the sequencing reads in order of genomic coordinates, to remove PCR duplicates, and to fix the mate information of the paired-end sequence read.

Somatic mutations were identified via local assembly of haplotypes using Mutect2.^23^Following GATK Best Practices recommendation, FilterMutectCalls GATK4 (version 4.0.8.0.1) was used for confident somatic calls, removed reads with base quality lower than 30, and filtered germline variants with gnomAD.^24^ We selected missense mutations with more than 20 depth and more than five alternative allele counts and manually reviewed calls with Integrative Genomics Viewer (IGV).

The patient-derived reliable somatic mutations and patient HLA genotyping information, and RNA sequencing data were prepared as input for the NoePepsee to predict patient-specific neoepitope candidates.^15^ NeoPepsee classifies potential neoantigens into three categories concerning predicted immunogenicity: high, medium, and low levels. Peptides at the high and medium levels were selected as candidates. pVACseq pipeline was run at the Carreno laboratory at the University of Pennsylvania as reported with a modified selection criteria (IC50 <100nM instead of 500nM, DNA VAF>20, and RNA VAF>0).^18^

### Processing of donor blood

Leukoreduction system (LRS) chambers from anonymous donors were obtained from the central bank. After removing 500 microliters for clinical HLA typing, a buffy coat was collected after ficoll gradient centrifugation at 2,000 rpm for 25 minutes at room temperature. Buffy coat was washed in PBS and resuspended in CellBanker2 solution (AMS Biotechnology Ltd, Abingdon, U.K.) and stored in the deep freezer.

### Clinical HLA genotyping of donor blood

DNA was extracted from 300 μL whole blood using the Gentra Puregene Blood Kit (QIAGEN, Hilden, Germany). Alternatively, DNA from PBMC was isolated using the DNeasy Blood & Tissue Kit (QIAGEN). DNA was eluted in 30 μL nuclease-free water, and samples were stored in a –80°C deep freezer until analysis. Sequences were analyzed with BIOWITHUS Inc. (Seoul, Korea).

### Peptide synthesis

All peptides were synthesized, and HPLC purified to over 98% by AnyGen (Gwangju, Korea) and was resolved in DMSO at a stock concentration of 10mg/ml. A working concentration of 10ug/ml was used for the ELISpot assay.

### ELISpot assay

White blood cells from HLA-matched donors were subject to perform an ELISpot assay. IFN-Ɣ pre-coated ELISpot assay kit (Cellular Technology Limited, Cleveland, Ohio) was used to determine the frequency of IFN-Ɣ generating cells.^17^ ELISpot assay was calibrated with influenza or CMV viral peptides.

Briefly, frozen PBMC isolated from LRS chambers were thawed and rested overnight in a culture medium consisting of CTS optimizer T cell expansion medium SFM (Gibco), human serum (5%), and 20 Unit Interleukin-2 (Gibco). PBMC were plated to U-bottom 96 well plate (10^5^cells per well) in culture medium. PBMC in each well was incubated with a corresponding peptide (10ug/ml) at three days intervals for peptide stimulation. After three cycles of peptide stimulation, stimulated PBMC were re-plated onto IFN-Ɣ coated ELISpot plate (2×10^5^ cells per well) followed by peptide re-stimulation for 24 hours. After re-stimulation with the peptide (on day 10), IFN-Ɣ production by the specific T-cell response to each peptide was estimated by ELISpot assay according to the manufacturer’s instruction. Peptide-stimulated PBMC was evaluated as compared with media (no-peptide) control, and phytohemagglutinin (PHA) was used as a positive control. ELISpot CTL reader was used for scanning ELISpot assay plate, and Immunospot software was used to analyze spot information.

Since there are multiple alleles expressed in each donor blood, we used three donors for each testing, and a peptide was called true neoepitope only when more than two among 3 HLA matched donor PBMCs showed spot counts more than 20% above no peptide control, upon rechallenge with the peptide after ten days incubation with the peptide and cytokines. PHA was used as a positive control.

## RESULT

### Patient characteristics

Among six primary breast cancer patients, 3 were clinically triple-negative subtype, a type of breast cancer with negative expression of estrogen, progesterone, and human epidermal growth factor receptor-2 (HER2), and 3 were hormone-receptor-positive breast cancer with luminal B feature, a subtype with aggressive clinical and biological features (Case ID, NEO; Table 1). In hormone-receptor-positive breast cancer patients, 2 showed invasive lobular carcinoma histology. At the time of sample collection, patients were not undergoing therapy except one patient (NEO2). NEO2 patient had received neoadjuvant chemotherapy (AC [Doxorubicin/ Cyclophosphamide]) followed by weekly paclitaxel regimen) but withdrew neoadjuvant chemotherapy after one paclitaxel infusion due to early disease progression and underwent surgery.

**Table 1.** Patient demographics.

| Case ID | Age | Histologic Type | Histologic Grade | Ki67 (%) | Sample source | Subtype | Stage | HLA type |
| --- | --- | --- | --- | --- | --- | --- | --- | --- |
| NEO1 | 60 | IDC | 3 | 77.2 | Surgery | TNBC | Stage IIA (pT1N1M0) | A*24:02, A*33:03 |
| NEO2 | 47 | IDC | 3 | 66.1 | Surgery | TNBC | ypStage IA (ypT1N0M0) | A*24:02, A*33:03 |
| NEO3 | 74 | IDC | 2 | 63.4 | Surgery | Luminal B BC | Stage IIA (pT2N0M0) | A*02:06, A*11:01 |
| NEO4 | 52 | ILC | 2 | 55.6 | Surgery | Luminal B BC | Stage IIA (pT2N0M0) | A*02:06, A*11:01 |
| NEO5 | 52 | ILC | 2 | 47.2 | Surgery | Luminal B BC | Stage IIA (pT2N0M0) | A*02:01, A*02:01 |
| NEO6 | 74 | IDC | 3 | 82.3 | Surgery | TNBC | Stage IIA (pT2N0M0) | A*02:01, A*24:02 |
| AV10 | NA | Squamous cell carcinoma | NA | NA | PDX | SQCC | stage IIB (T2bN1M0) | A*32:01, A*33:03 |
| AV13 | NA | Non-small cell | NA | NA | PDX | NSCLC | stage IIIA (T2bN2 M) | A*02:01, A*30:04 |
| AV38 | NA | Squamous cell carcinoma | NA | NA | PDX | SQCC | stage 1b | A*01:01, A*31:01 |

IDC; invasive ductal carcinoma, ILC; invasive lobular carcinoma, TNBC; triple-negative breast cancer, BC; breast cancer, SQCC; squamous cell carcinoma, NSCLC; non-small cell carcinoma of the lung, PDX; patient-derived tumor xenograft

Two PDX established from surgical specimens from squamous cell carcinoma of the lung (AV10 and AV38) and one from non-small cell lung cancer (AV10) were used to represent lung cancer (Table 1).

### Comparison of HLA genotyping methods

We used Sanger sequencing-based clinical HLA genotyping results with two-digit resolution (BIOWITHUS Inc. Seoul, Korea). In addition, we applied Optitype ^19^ and HLAminer ^20^and used the Omixon Holotype HLA™ kit (Omixon Biocomputing Ltd, Budapest, Hungary) to evaluate NGS-based HLA typing methods (Table 2).

**Table 2.** HLA genotyping results comparison among three different methods.

| Case ID | HLA-A genotype<br>(clinical) | OptiType | HLAminer | Omixon Holotype |
| --- | --- | --- | --- | --- |
| NEO1 | A*24:02, A*33:03 | A*24:02, A*33:03 | A*24:02, A*33:03 | A*24:02, A*33:03 |
| NEO2 | A*24:02, A*33:03 | A*24:02, A*33:03 | A*24:02, A*33:03 | A*24:02, A*33:03 |
| NEO3 | A*02:06, A*11:01 | A*02:06, A*11:01 | A*02:06, A*11:01 | A*02:06, A*11:01 |
| NEO4 | A*02:06, A*11:01 | A*02:06, A*11:01 | A*02:06, A*11:01 | A*02:06, A*11:01 |
| NEO5 | A*02:01, A*02:01 | A*02:01, A*02:01 | A*02:01, A*02:01 | A*02:01, A*02:01 |
| NEO6 | A*02:01, A*24:02 | A*02:01, A*24:02 | **A\*24:95, A\*69:01** | A*02:01, A*24:02 |
| AV10 | A*32:01, A*33:03 | **A\*33:03**, A*33:03 | **A\*33:147, NA** | ND |
| AV13 | A*02:01, A*30:04 | A*02:01, A*30:04 | **A\*30:04, A\*69:01** | ND |
| AV38 | A*01:01, A*31:01 | A*01:01, A*31:01 | A*01:01, **A\*36:03** | ND |
| Average<br>accuracy | reference | 17/18 (94%) | 11/18 (61%) | 12/12 (100%) |

Note: bold words represent mismatches

There was a high concordance between NGS-based Optitype and clinical HLA genotyping tests with 94% agreement (Supplementary Table 1). However, the concordance rate for HLAminer was low (61%). Omixon Holotype assay results were 100% concordant with clinical genotypes.

These results suggest that HLA typing with OptiType algorithm for exome or RNAseq data are good alternatives for clinical HLA typing tests when analyzing candidate neoepitopes selection, without the need to use costly dedicated kits for HLA genotyping.

### Candidate neoepitope selection

Paired whole-exome sequencing of tumor and PBMC DNA yielded an average of 95 (range 70 to 107) nonsynonymous somatic mutations in breast cancer tumors and an average of 109 (range 76 to 143) nonsynonymous mutations in lung cancer PDX samples, respectively.

In this study, we focused only on HLA-A allele restricted neoepitopes for experimental validation. Neopepsee considers only SNVs and does not consider fusion genes or intron retention. Neopepsee picked an average of 19 candidates (range 7 to 33) from breast cancer mutations and 17 candidates (range 11 to 21) from lung cancer mutations.

For pVACseq analysis with a modified threshold of IC50 below 100 nM, DNA VAF >20 and RNA VAF >0 yielded an average of 9 best candidates for breast cancer (range 3 to 26) and 15 for lung cancer PDX samples (range 8 to 21). In total, Neopepsee selected 159 out of 898 mutations, and pVACseq selected 84 mutations with only 26 mutations shared between the two prediction algorithms.

### *In vitro* validation of candidate neoepitopes

Based on a report by Stronen et al., we used HLA matched donor blood for ELISpot assay of IFN-Ɣ secretion from the neoantigen-specific T cell populations to validate candidate neoepitopes. Twenty-two Neopepsee candidate peptides and nine pVACseq candidate peptides could not be tested either due to synthesis or purification failure or due to lack of donor blood for the specific HLA alleles. The results are summarized in Table 3. In total, 36 of 191 tested candidates (18.8%) were positive by ELISpot.

**Table 3.** Summary of ELISpot assays of IFN-Ɣ secretion from neoantigen-specific T cell populations from donor PMBCs.

|  | HLA genotype | N of Mutations | NeoPepsee |  |  |  | pVACseq (modified)* |  |  |  | Common to both |  |  |  |
| --- | --- | --- | --- | --- | --- | --- | --- | --- | --- | --- | --- | --- | --- | --- |
|  |  |  | predicted | tested | ELISpot positive | PPV | predicted | tested | ELISPO T positive | PPV | predicted | tested | ELISpot positive | PPV |
| NEO 1 | A*24:02, A*33:03 | 70 | 9 | 9 | 3 | 0.33 | 4 | 4 | 1 | 0.25 | 3 | 3 | 1 | 0.33 |
| NEO 2 | A*24:02, A*33:03 | 94 | 7 | 6 | 2 | 0.33 | 7 | 6 | 0 | 0.00 | 1 | 0 | 0 |  |
| NEO 3 | A*02:06, A*11:01 | 103 | 28 | 27 | 6 | 0.22 | 26 | 25 | 4 | 0.16 | 11 | 10 | 2 | 0.20 |
| NEO 4 | A*02:06, A*11:01 | 96 | 33 | 31 | 1 | 0.03 | 3 | 3 | 0 | 0.00 | 1 | 1 | 0 | 0.00 |
| NEO 5 | A*02:01, A*02:01 | 100 | 12 | 12 | 4 | 0.33 | 5 | 5 | 2 | 0.40 | 2 | 2 | 1 | 0.50 |
| NEO 6 | A*02:01, A*24:02 | 107 | 24 | 22 | 6 | 0.27 | 9 | 7 | 2 | 0.29 | 4 | 3 | 1 | 0.33 |
| AV10 | A*32:01, A*33:03 | 143 | 11 | 3 | 0 | 0.00 | 5 | 3 | 0 | 0.00 | 0 | 0 | 0 |  |
| AV13 | A*02:01, A*30:04<sup>#</sup> | 109 | 21 | 17 | 6 | 0.35 | 15 | 13 | 5 | 0.38 | 4 | 2 | 2 | 1.00 |
| AV38 | A*01:01<sup>#</sup>, A*31:01 | 76 | 14 | 10 | 0 | 0.00 | 10 | 9 | 1 | 0.11 | 0 | 0 | 0 |  |
| Total |  | 898 | 159 | 137 | 28 | 0.20 | 84 | 75 | 15 | 0.20 | 26 | 21 | 7 | 0.33 |

<sup>#</sup>Note: A\*30:04 and A\*01:01 matched peptides were not tested due to lack of donor blood

The median of four (range 1 to 6) NeoPepsee selected candidate neoepitopes from breast cancer cases, and the median of two (range 0 to 6) candidates from lung cancer cases were positive with ELISpot (Table 4). In comparison, the median of two (range 0 to 4) and the median of two (range 0 to 5) from pVACseq selected candidates from breast and lung cancer were positive with ELISpot, respectively. Positive predictive value (PPV) ranged from 0 to 0.35 for Neopepsee and 0 to 0.38 for pVACseq (Table 5). In total, 28 of 137 tested Neopepsee candidates were ELISpot positive (PPV 0.20) compared to 15 of 75 tested pVACseq candidates (PPV 0.20). Among 20 tested candidates predicted by both algorithms, seven were positive by ELISpot (PPV 0.35). Detailed raw data are provided in Supplementary Table 2. For each tumor sample, candidates by Neopepsee and pVACseq are listed, HLA genotypes of donor blood used for ELISpot, and results for each donor blood are shown in Table 4.

**Table 4.**
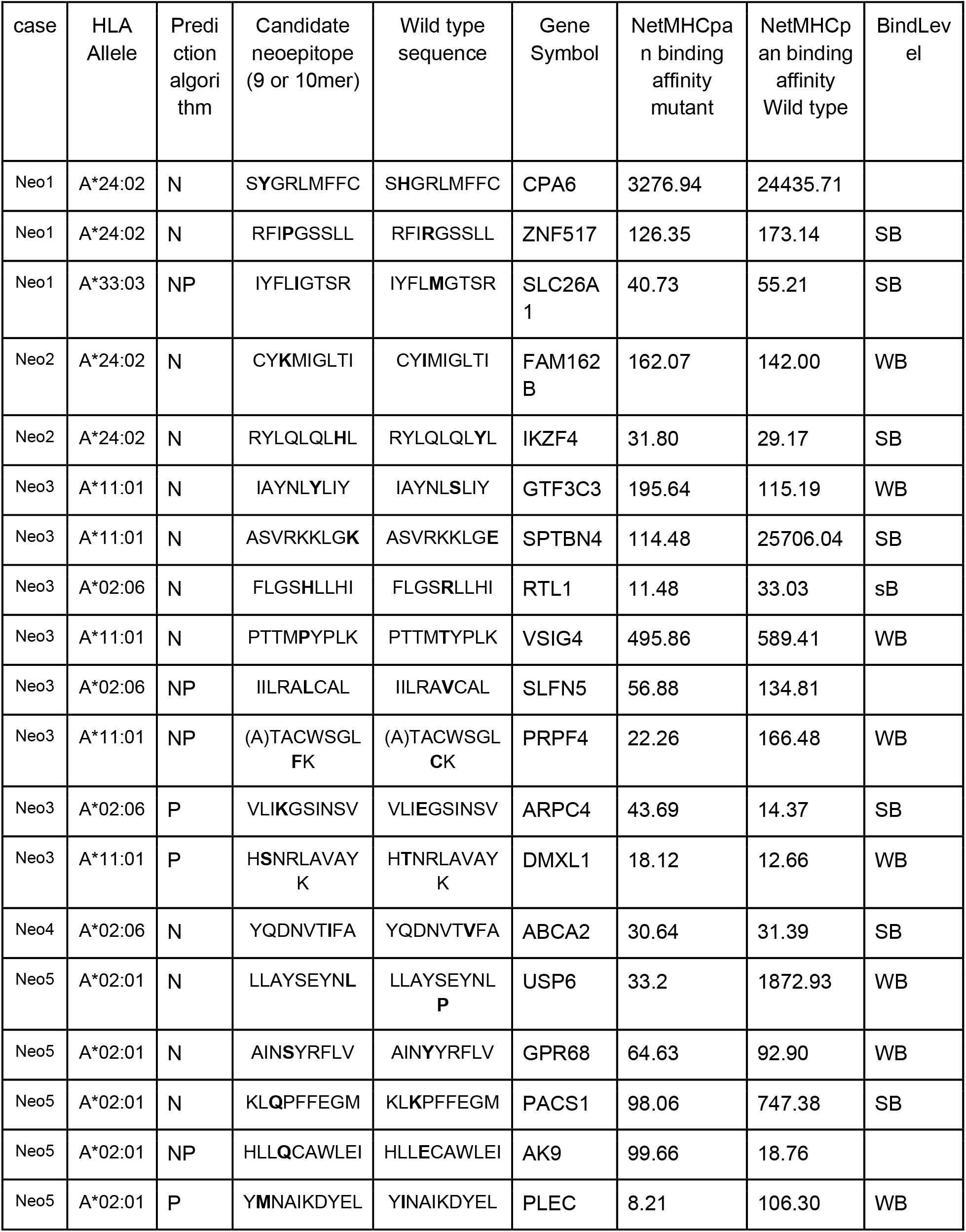
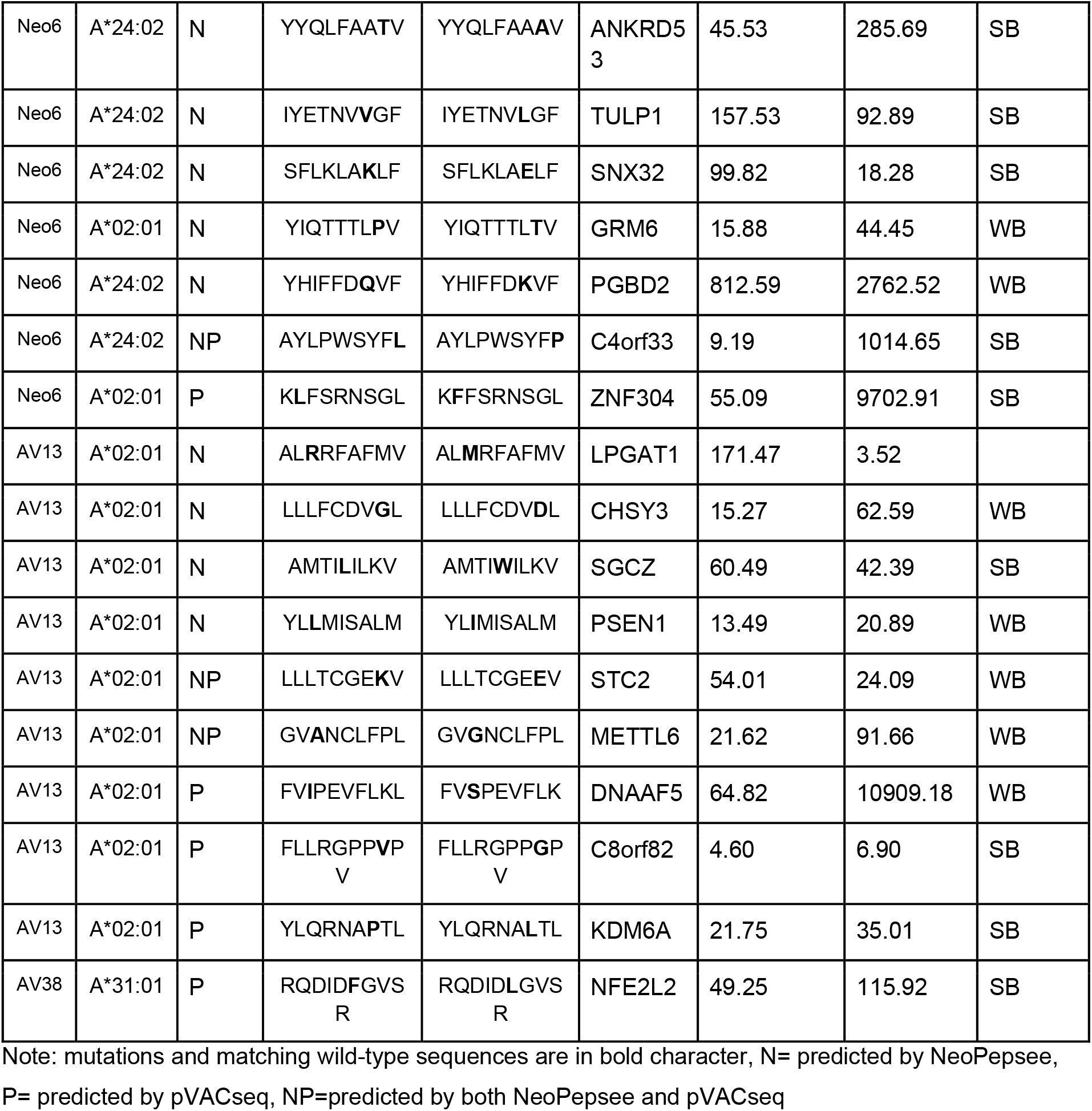
List of ELISpot-positive cancer-specific neoepitopes.

**Table 5.** Summary of *in vitro* validated neopeptides by ELIspot assay according to MHC binding affinity of 34 nM.

|  | IC<sub>50</sub> <34nM | IC<sub>50</sub> ≥34nM | Marginal row total |
| --- | --- | --- | --- |
| ELISpot positive | 14 (38.9%) | 22 (61.1%) | 36 |
| ELISpot negative | 47 (30.3%) | 108 (69.7%) | 155 |
| Marginal column total | 61 (31.9%) | 130 (68.1%) | 191 |

For some candidate peptides, although no memory response could not be demonstrated, there were high spots compared to the wells not stimulated with the peptide from day 0. We did not consider those peptides as positive. For some peptides, there were strong responses in only one of the three donor blood, suggesting that the response could be directed toward the other allele than the predicted one.

### Evaluation of TESLA recommended criteria for neoepitope in Yonsei cohort

Previously, the Tumor Neoantigen Selection Alliance (TESLA) suggested that potential immunogenic peptides are characteristic of MHC binding affinity stronger than 34 nM^14^.

According to these criteria, we compared the validated neopeptides based on MHC binding affinity of 34 nM (Table 5).

Fourteen of 36 (38.9%) ELISpot positive peptides had NetMHCpan IC_50_ below 34nM, whereas 47 of 155 (30.3%) ELISpot negative peptides had IC^50^ below 34nm (Fisher’s exact test, *p*=0.32, NS). If we were to predict based on IC_50_ value, 61/191 (31.9%) peptides are predicted to be high-affinity binders, and among them, 14 (22.9%) are true positives. Therefore, we could not corroborate the findings from the TESLA consortium.

## DISCUSSION

Clinical trials for personalized therapeutic neoepitope vaccines have demonstrated that T-cell immune responses were induced for only a subset of candidate neoepitopes.^8–11^Although almost all vaccine trials reported so far selected candidates based on computational algorithms, one trial selected candidates based on *in vitro* T-cell response to candidate neoepitopes presented by bacteria.^13^ Intriguingly, in the latter study, some mutated peptides suppressed immune response and investigators named such peptides as ‘inhibigens’. Inhibigens suppressed immune activation by otherwise immune-stimulatory neoepitopes when injected together. Since *in vitro* validation of every candidate neoepitope is demanding and time-consuming, it is challenging to incorporate into a routine process. Thus, designing personalized therapeutic vaccines requires a robust computational neoepitope prediction algorithm with a high positive predictive value.

In 9 cases prospectively screened using ELISpot assay with HLA matched donor PBMC, PPV was 20% for both Neopepsee and pVACseq. There was relatively little overlap between the predicted epitopes between the two algorithms. PPV increased to 33% for overlapped candidates between Neopepsee and pVACseq. This finding is consistent with the TESLA report in that ensemble models outperform individual prediction models.

TESLA identified a set of thresholds for several variables that in combination could filter out 93% of non-immunogenic peptides while maintaining 55% of immunogenic peptides.^14^ This threshold set is composed of binding affinity less than 34 nM, tumor abundance greater than 33 TPM, and binding stability greater than 1.4 h. We examined the utility of this threshold in our cases. Fourteen of 36 (38.9%) ELISPOT positive peptides had NetMHCpan IC_50_ below 34 nM, whereas 47 of 155 (30.3%) ELISPOT negative peptides had binding affinity below 34 nM (Fisher’s exact test, *P*=0.32). If we were to predict based on the binding affinity threshold suggested by the TESLA, 61/191 (31.9%) peptides are predicted to be high-affinity binders, and among them, 14 (22.9%) are true positives. Therefore, we could not corroborate the findings from the TESLA consortium. Unlike TESLA, which used a multimer assay to validate the neoepitopes, we used an ELISpot assay. While a multimer assay quantifies T-cells with T-cell Receptors that can bind MHC presented neoepitopes, ELISpot assay quantifies the Interferon-Ɣ secreted by the T-cells activated by the MHC presented neoepitopes. The differences in sensitivity and specificity between the two assays may be partially responsible for the observed difference with the TESLA data.

Our study has several limitations: 1) We did not screen mutated peptides that were not predicted by the Neopepsee or pVACseq. Since there is a possibility that some of them might be immunogenic, we do not know the true false-negative rate of each algorithm. 2) Due to the limitation of the amount the blood drawn from patients, we used HLA-matched donor blood for the ELISpot assay. Stronen et al. reported that HLA-matched healthy donor blood showed a five times higher response rate than blood from patients. Thus our result of 20% PPV could be an overestimate.^16^ 3) For ELISpot assay, we used strict criteria of requiring memory response upon rechallenge with the candidate peptides after ten days incubation with the peptides. Some of the peptides showed increased spot numbers over control wells that have never contacted the peptides but did not show increased spot number in comparison to no peptide control at the time of rechallenge. Such results could be interpreted as positive responses by some investigators. 4) ELISpot is known to be influenced by many factors.^25^ Compared to viral antigens, which show strong positive ELISpot results, most of the neoepitopes we found showed a low to moderate increase in spot numbers. Since we did not validate our results with a multimer assay, some peptides were presumably false-positive findings.

In conclusion, our data replicate the findings from many studies that the current generation of neoepitope prediction algorithms suffer from low PPV. This is most likely due to a paucity of positive T-cell immune response data in the training database. In a training dataset used for a recent effort to develop a prediction algorithm, only 129 of 3329 cancer mutations were experimentally tested positive neoepitopes.^26^ Thus, more publicly available *in vitro* validation data is required to train the prediction algorithms to achieve clinically meaningful PPV. Our data will add a significant new information to the publicly available database of neoepitopes.

---

**Supplementary table 1.**
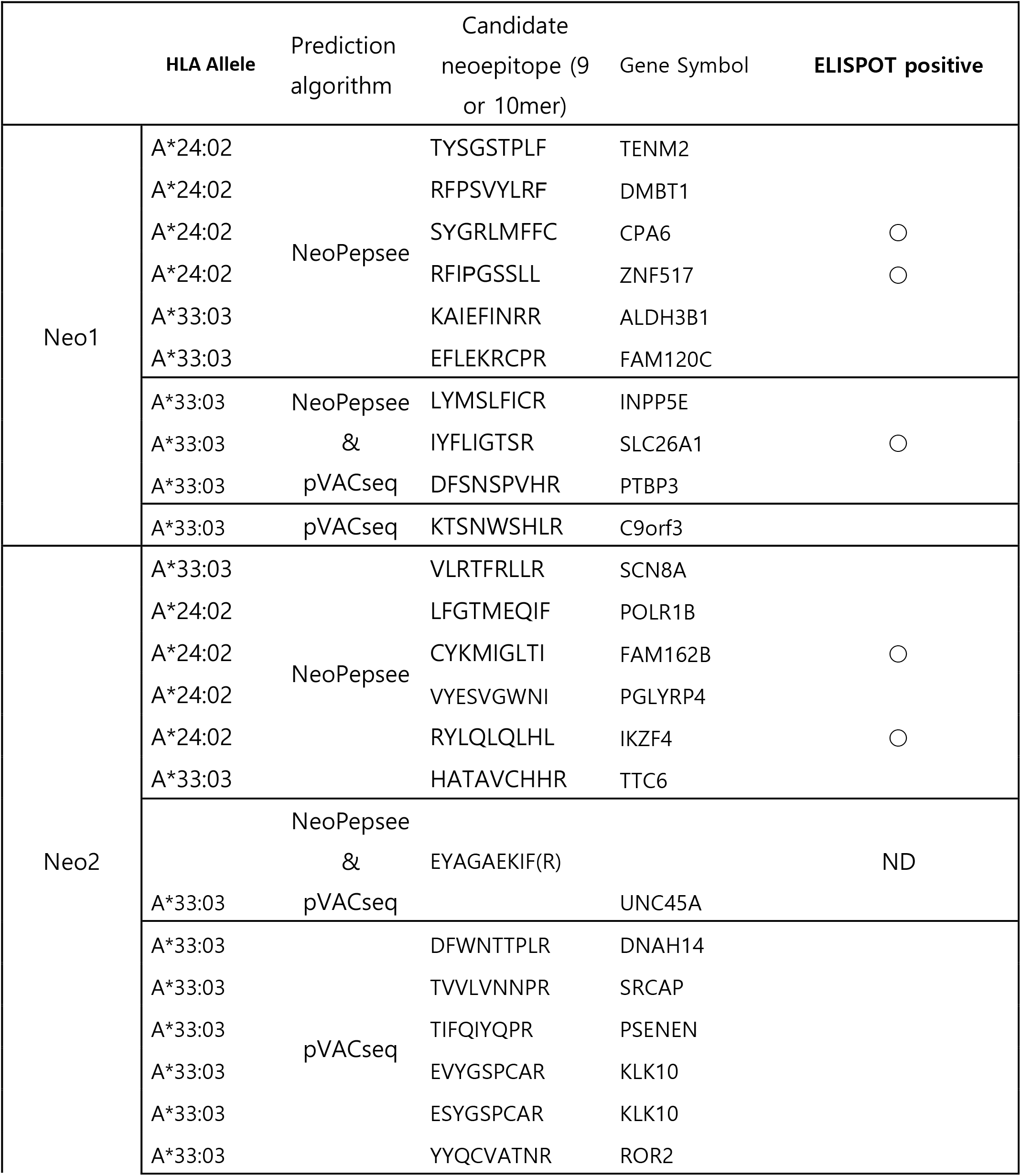
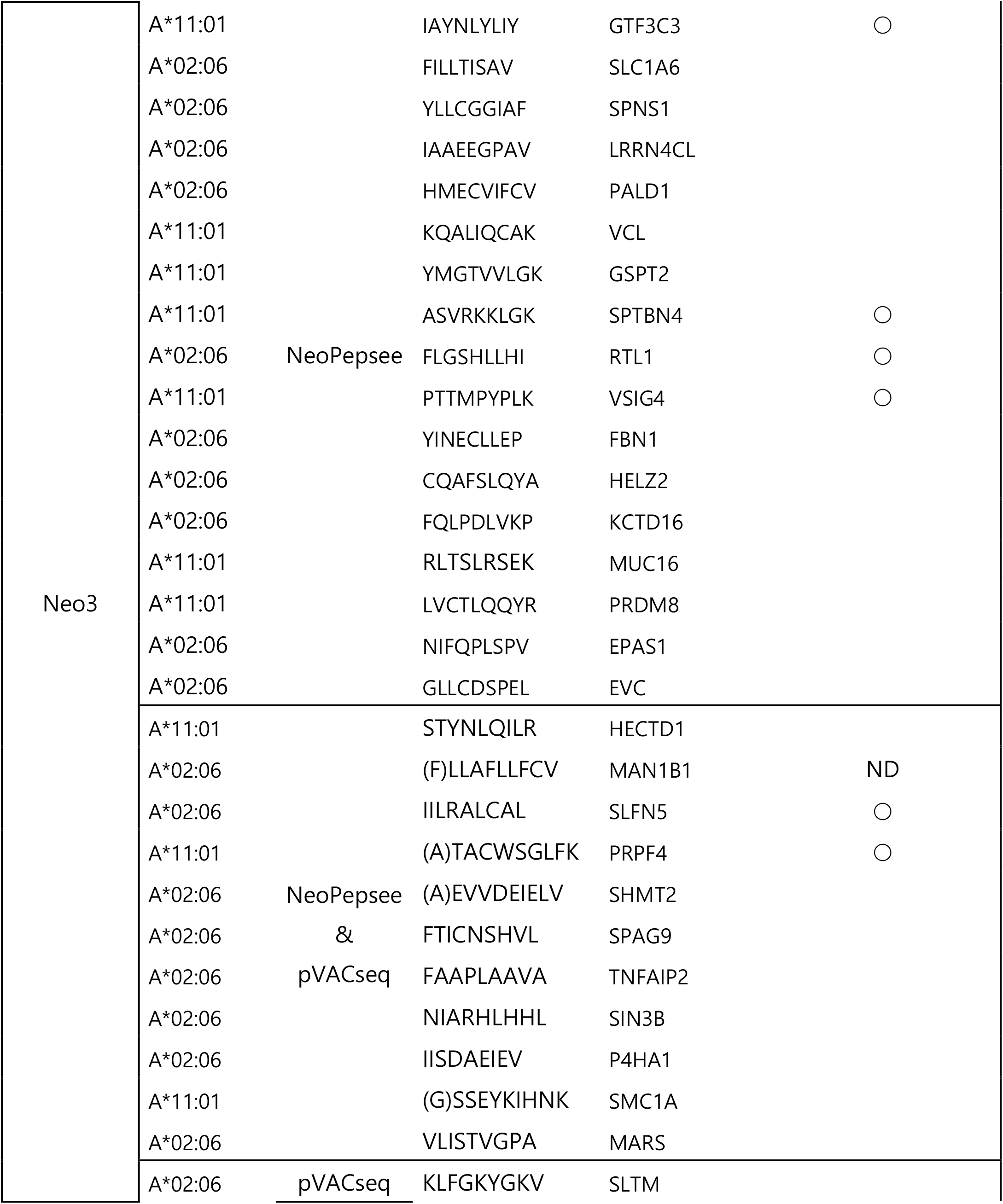
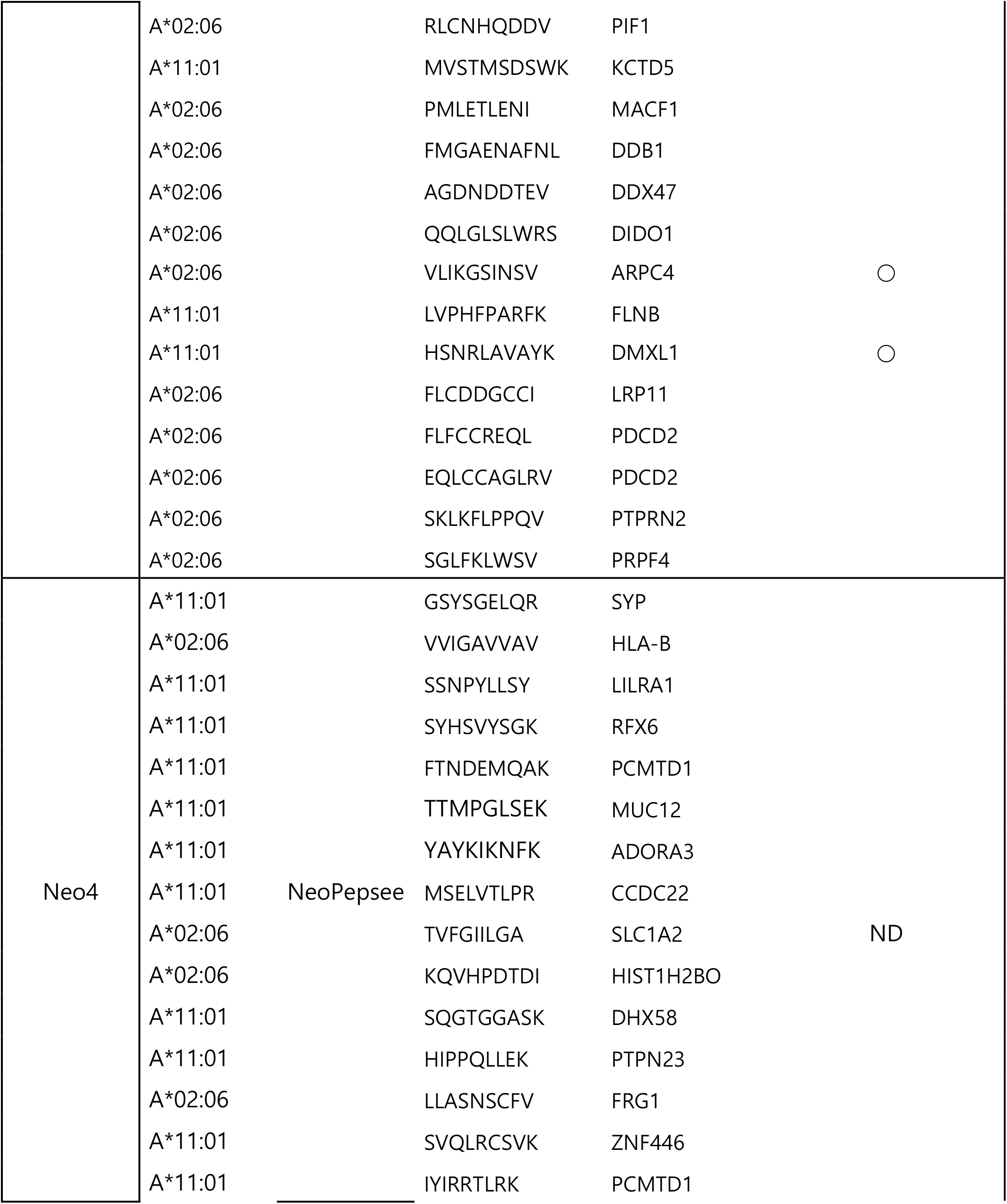
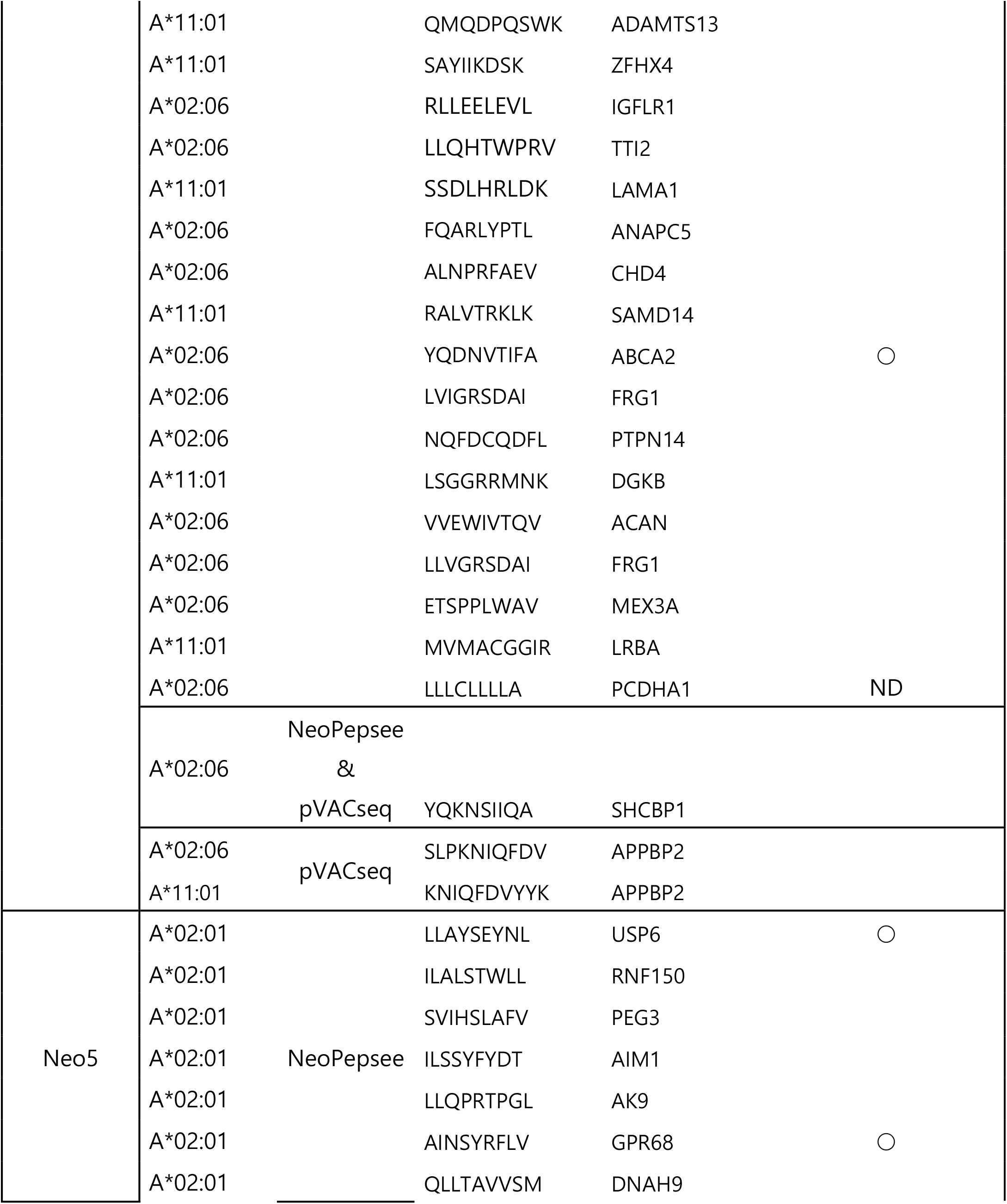
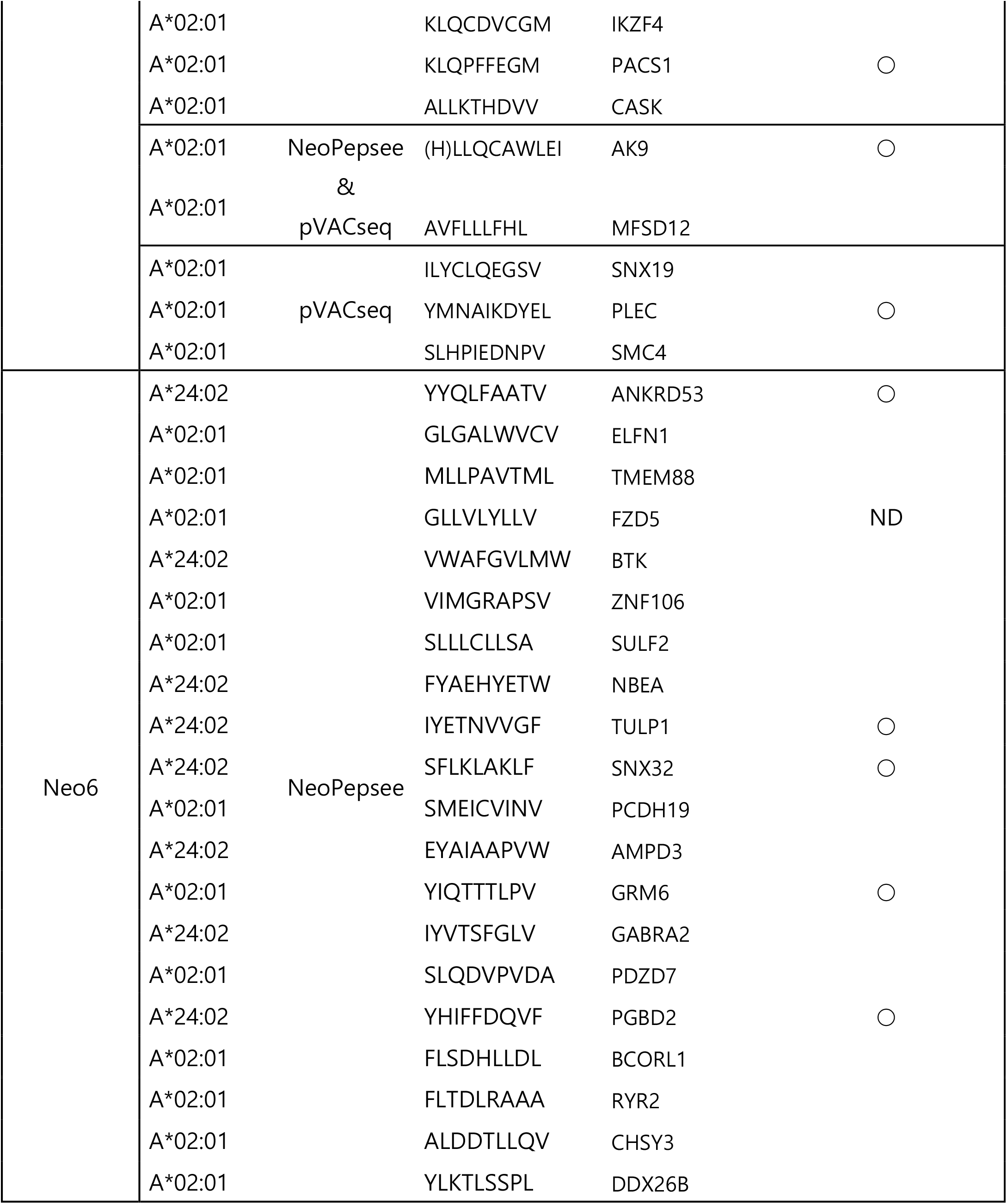
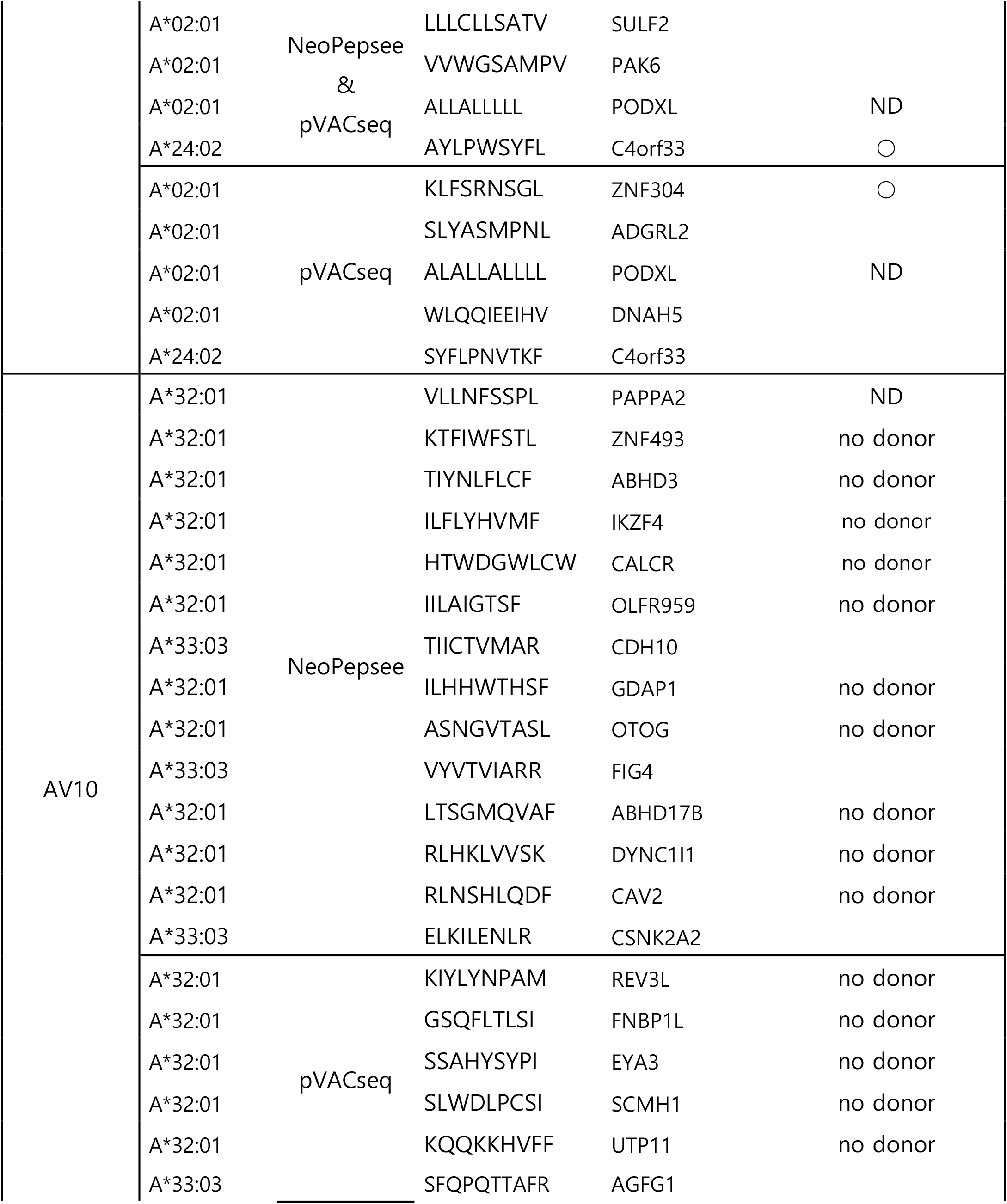
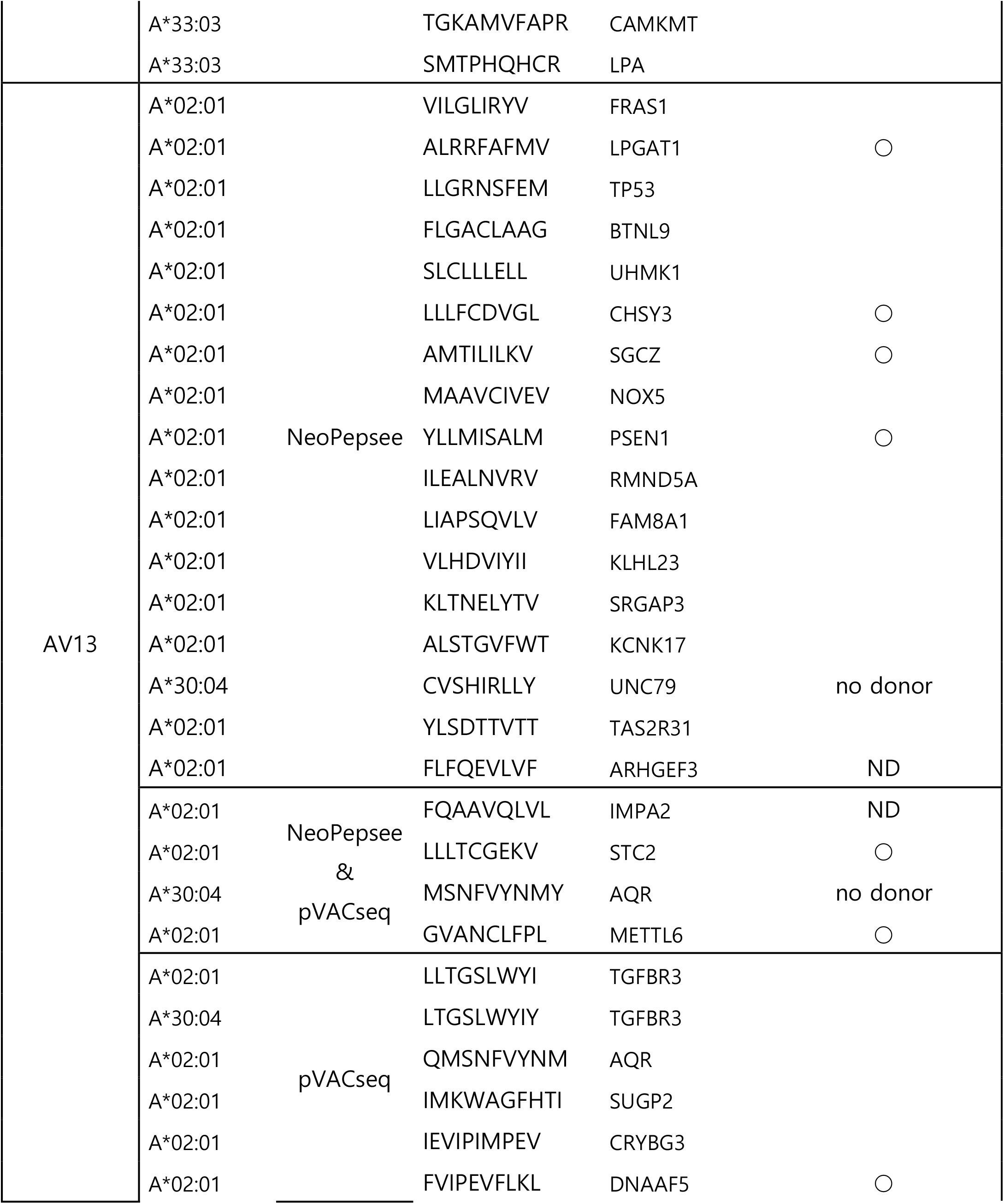
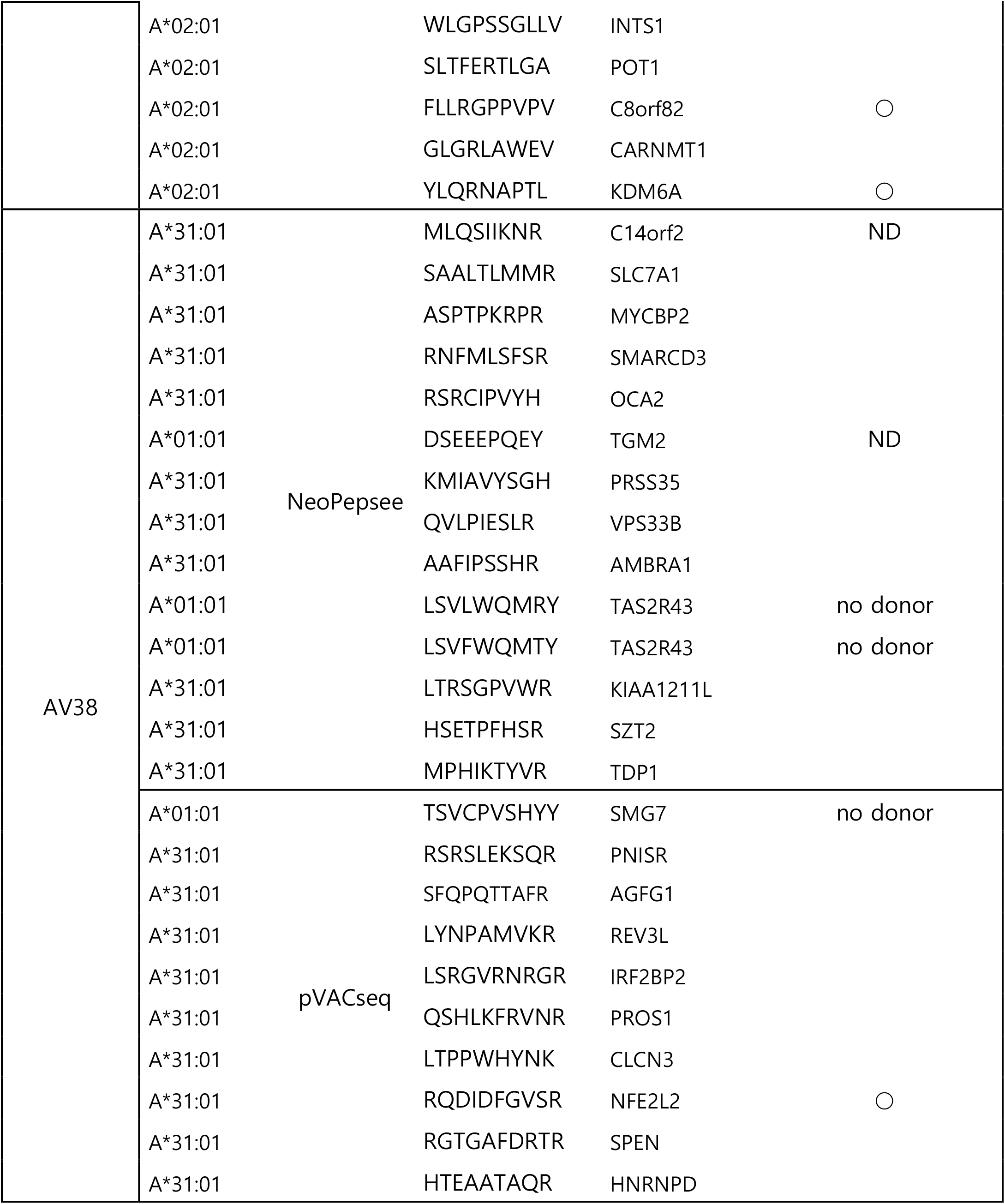
List of candidate peptides

